# Immunoglobulin heavy chains are sufficient to determine most B cell clonal relationships^1^

**DOI:** 10.1101/665760

**Authors:** Julian Q. Zhou, Steven H. Kleinstein

## Abstract

B cell clonal expansion is vital for adaptive immunity. High-throughput B cell receptor (BCR) sequencing enables investigating this process, but requires computational inference to identify clonal relationships. This inference usually relies on only the BCR heavy chain, as most current protocols do not preserve heavy:light chain pairing. The extent to which paired light chains aids inference is unknown. Using human single-cell paired BCR datasets, we assessed the ability of heavy chain-based clonal clustering to identify clones. Of the expanded clones identified, <20% grouped cells expressing inconsistent light chains. Heavy chains from these misclustered clones contained more distant junction sequences and shared fewer V segment mutations than the accurate clones. This suggests that additional heavy chain information could be leveraged to refine clonal relationships. Conversely, light chains were insufficient to refine heavy chain-based clonal clusters. Overall, the BCR heavy chain alone is sufficient to identify clonal relationships with confidence.

## Introduction

B cell-mediated immunity relies on immunoglobulin (Ig) antibodies produced as a result of B cell clonal expansion. A B cell receptor (BCR) is the membrane-bound form of an antibody, and is made up of heavy and light chains paired in a heterodimeric fashion. Each chain contains a variable (V) region, and together the V regions from the heavy and light chains form the antigen-binding sites. The V regions are formed via V(D)J recombination. In human, this shuffling process brings together one gene each from numerous IGHV, IGHD, and IGHJ genes for the heavy chain V (VH) region; and one gene each from either IGKV and IGKJ genes, or IGLV and IGLJ genes for, respectively, the κ or the *λ* light chain V (VL) region. Enzyme-mediated editing of the V(D)J junctions and the pairing of heavy and light chains inject additional diversity (1). During adaptive immune responses, B cells proliferate and further diversify via somatic hypermutation (SHM), forming clones consisting of cells which originated from the same V(D)J recombinant events, yet whose BCRs differ at the nucleotide level. As a result, each BCR is largely unique, with recent estimate suggesting 10^16^-10^18^ unique paired antibodies in the circulating repertoire (2).

Adaptive Immune Repertoire Receptor sequencing (AIRR-seq) allows for high-throughput profiling of the diverse BCR repertoire via full-length V(D)J sequencing in bulk (3). An ensuing challenge is to computationally infer B cell clonal relationships (4). This step is of great importance as the assessment of repertoire properties such as diversity (5) depends on proper identification of clones, as does the reconstruction of B cell clonal lineage (6) for tracing isotype switching (7) and antigen-specific (8) antibodies. To infer clones, differences at the sequence nucleotide level, especially the high diversity in the CDR3 region, can serve as “fingerprints” (9). Likelihood-based (10) and distance-based (11–14) approaches exist. For instance, cells sharing the same IGHV and IGHJ genes, and whose heavy chain junctional sequences are sufficiently similar based on a fixed (11–13) or adaptive (14) distance threshold, may be clustered as clones. For validation, existing methods used simulated and experimental heavy chain sequences (10, 13, 14), measuring the fractions of sequences inferred to be clonally unrelated and related of being, respectively, truly unrelated and related (specificity and sensitivity). Recently, Nouri & Kleinstein reported both metrics at over 96% based on simulated data (14).

The majority of current BCR repertoire studies utilizes bulk sequencing (15), during which VH:VL pairing is lost (16). In the absence of VH:VL pairing, computational methods for identifying clones have focused on heavy chain BCR data. This is justified under the assumption that heavy chain junctional diversity alone should be sufficiently high such that, even without light chains, the likelihood of clonally unrelated cells being clustered together will be negligibly small (13). This reasoning has yet to be rigorously tested with experimental data. Recent breakthroughs in single-cell BCR sequencing technology have enabled the recovery of native VH:VL pairing (17, 18). We now have the opportunity to investigate the extent to which inclusion of light chains impacts the ability to accurately detect B cell clonal relationships.

Using single-cell VH:VL paired BCR data, we assessed the performance of heavy chain-based computational methods for identifying clones by measuring the extent to which the inferred clonal members expressed consistent light chains sharing the same V and J genes and junction length. We conclude that clonal members of the majority of the inferred clones exhibited light chain consistency. For the majority of the accurately inferred heavy chain-based clones, light chain information did not lead to further clonal clustering with greater granularity. At least some of the information gained from paired light chain data was apparent when considering the pattern of shared mutations in the heavy chain V segment, which is not considered in current distance-based clonal clustering methods, thus offering the potential for further improvements in heavy chain-based clonal inference.

## Materials and Methods

### Single cell immune profiling datasets

Four human datasets (**Supplemental Fig. 1A**) published by 10x Genomics for public use on 8/1/2018 were accessed (https://support.10xgenomics.com/single-cell-vdj/datasets) on 11/3/2018. Two datasets were sorted and produced by direct Ig enrichment of, respectively, CD19+ B cells isolated from PBMCs from a healthy donor, and GM12878 B-lymphoblastoid cell line. They contain VH:VL paired reads for individual cells. The other two datasets were unsorted and produced by V(D)J+5’ gene expression profiling of, respectively, PBMCs from a healthy donor, and squamous non-small cell lung carcinoma (NSCLC) cells from a fresh surgical resection. These contain gene expression measurements and Ig enrichment with VH:VL pairing. All datasets were outputted by 10x Genomics via Cell Ranger (v2.2.0). We used “filtered contigs” and “filtered gene-barcode matrices” for Ig and gene expression respectively.

A fifth dataset (**Supplemental Fig. 1B**) from Croote et al. contains BCR contigs covering full-length V(D)J segments reconstructed from single-cell RNA-seq of FACS-sorted CD19+ B cells from six food-allergic individuals (19).

### Germline V(D)J gene annotation of BCR contigs

Germline V(D)J gene annotation was performed using IMGT/HighV-QUEST and IgBLAST (v1.10.0). The germline reference used was IMGT release 201839-3. The 10x Genomics datasets also contained annotations by Cell Ranger. IMGT/HighV-QUEST annotations were used as final annotations post-filtering.

### Filtering of BCR contigs and cells

For all datasets, only productively-rearranged BCR contigs with valid V and J gene annotations, consistent chain annotation (excluding such contigs with IGHV and IGK/LJ), and junctions with nucleotide lengths being a multiple of 3 were used. A contig must meet all abovementioned criteria based on annotations from all programs used. Furthermore, only cells with exactly one heavy chain contig paired with at least one light chain contig were examined. From the two unsorted 10x Genomics datasets with gene expression, we considered only cells displaying a transcriptomic profile consistent with being a B cell. Taking into account of the high dropout rate of single cell RNA-seq, B cells were defined as any cell with non-zero log-normalized expression for any one of these genes: pan-B cell markers CD19, CD24, and CD72 (20), plasmablast markers CD38 and MKI67 (21), and the isotype-encoding genes.

### Heavy chain-based B cell clonal clustering

For each dataset, on a per-subject basis, we identified clones using distance-based methods. We used, separately, spectral clustering (SCOPer v0.1.999) (14) and hierarchical clustering (13) (Change-O v0.4.3) (22). Both methods first partitioned cells into groups sharing the same combination of IGHV gene, IGHJ gene, and heavy chain junction length (heavy chain VJL combination), where junction is defined as the IMGT-numbered codon 104 (conserved Cys) to codon 118 (conserved Phe/Trp) (23). Within each group, based on distances among the heavy chain junction sequences, a threshold was used to cluster cells within that group into clones. For spectral clustering, adaptive thresholds were chosen by an unsupervised machine learning algorithm. For hierarchically clustering, a subject-specific, fixed threshold was chosen upon inspection of distance-to-nearest-neighbor plots (**Supplemental Fig. 1C**) (13).

### Calculation of the number and frequency of nucleotide mutations

The number and frequency of nucleotide mutations were calculated based on IGH/K/LV positions leading up to the junction region using the “calcObservedMutations” function from SHazaM (v0.1.10) (22). To calculate the number of IGHV mutations shared pairwise between cells from the same clone, we counted the number of positions at which mutations involving the same nucleotide change were observed in both cells.

## Results

We performed clonal relationship inference for five single-cell, VH:VL paired, human BCR datasets, using only the heavy chain sequence from each cell. The datasets included four publicly available ones from 10x Genomics and one described by Croote et al. (19) (**Materials & Methods**). Of these, the B-lymphoblastoid GM12878 cell line dataset served as positive control, as any clone present in a cell line culture can be expected to comprise genetically identical clonal members. A distance-based spectral clustering method (14) was applied to identify clones for each dataset. The datasets contained between 3 and 157 non-singleton clones (i.e., clones containing at least two cells) (**Table 1**). Due to the small number of clones in each of the six food-allergic individuals in the Croote et al. dataset, we aggregated those results for display after performing analysis on a per-subject basis.

**Table 1:** Numbers of heavy chain-based clones and clone sizes.

| Dataset | Total<br>number<br>of clones | Number of<br>non-singleton<br>clones | Clone size of<br>non-singleton clones<br>(number of clones) |
| --- | --- | --- | --- |
| CD19+ B cells | 8268 | 157 | 2 (136); 3(15); 4 (4); 5 (2) |
| B cells from NSCLC tumor | 1247 | 98 | 2 (81); 3 (9); 4 (4);<br>6 (2); 8 (1); 14 (1) |
| B cells from food-allergic individuals | 952 | 12 | 2 (10); 3 (1); 5 (1) |
| B cells from healthy PBMCs | 1105 | 8 | 2 (6); 3 (2) |
| GM12878 cell line | 5 | 3 | 2 (1); 5 (1); 790 (1) |

### Heavy chain-based clonal clustering is accurate for over 80% of clones

To assess the extent to which heavy chain-based clonal clustering captures the underlying biological truth of B cell clonal relationships, we examined whether cells clustered into the same clone based on their heavy chains alone expressed consistent light chains. Specifically, within the same clone, cells that are truly clonally related should carry light chains comprised of the same combination of IGK/LV gene, IGK/LJ gene, and junction sequences of identical lengths (hereafter referred to as the V-J-junction-length, or VJL, combination). An inferred clone was considered accurate if all of its clonal members carried light chains with the same VJL combination, and “misclustered” otherwise. Using spectral clustering with adaptive thresholds, 83% to 97% of the inferred clones were accurate (**Figure 1A**). Another distance-based hierarchical clustering method using a fixed distance threshold (13) yielded similar results (**Figure 1B**), and therefore we focus on presenting results from spectral clustering hereafter. To test the possibility that the observed accuracy arose by chance, we randomly permuted the VH:VL pairings of the cells, while maintaining their heavy chain-based clustering structures. Across 100 permutations, only 1% to 6% (SD 1% to 8%) of the inferred clones were accurate by chance (**Figure 1A**). Overall, these results show that heavy chain-based clonal clustering can determine clonal relationships with reasonable confidence (>80%) in terms of light chain consistency.

**Figure 1.**
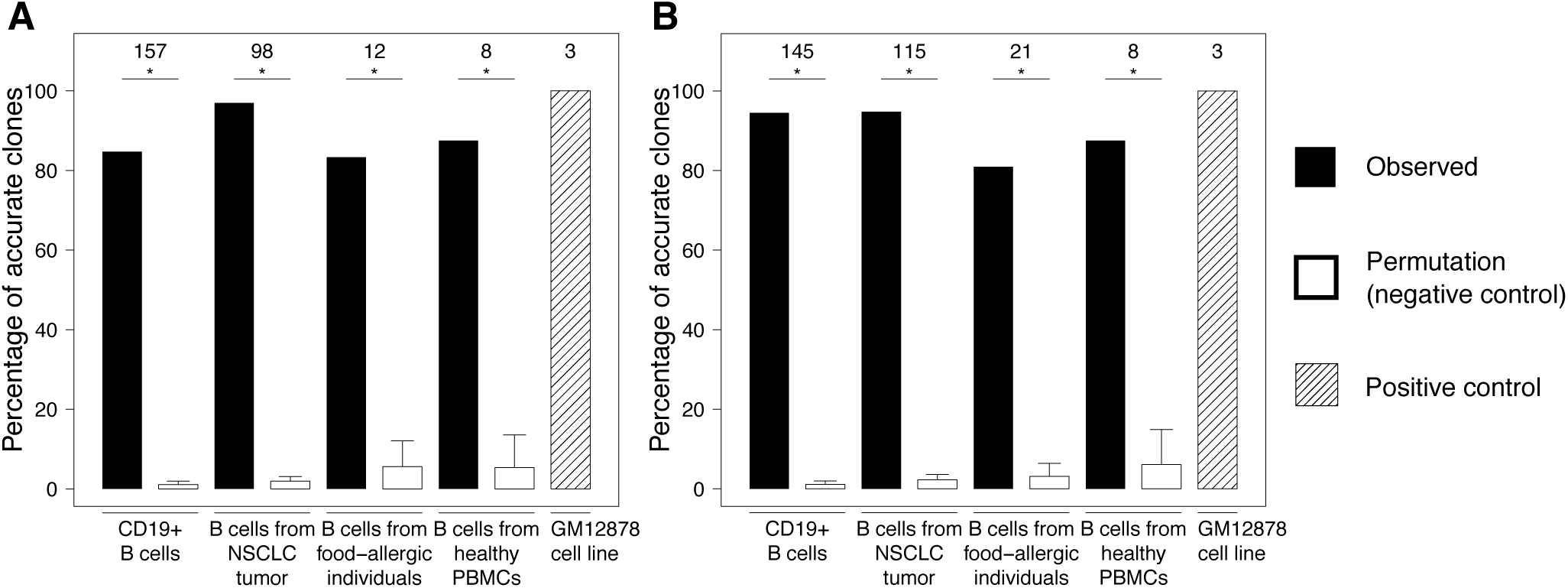
Performance of heavy chain-based **(A)** spectral clustering with adaptive threshold and **(B)** hierarchical clustering with fixed threshold. Solid and shaded bars show percentages of non-singleton inferred clones in which clonal members all carried light chains with the same V-J-junction-length (VJL) combination. Numbers at the top indicate the denominators. A background distribution was generated by permuting VH:VL pairings 100 times while maintaining inferred clonal lineage structures. Hollow bars show average percentages of accurate clones across permutations with standard errors. * denotes empirical one-sided, Bonferroni-corrected p-value < 0.05.

We next investigated the possibility that the observed level of confidence was deflated by factors unrelated to the clonal clustering method itself. We considered the possibility that the misclustered clones (**Supplemental Fig. 2A**) arose due to erroneous barcoding during sequencing preparation, which has the potential to link together heavy and light chains from unrelated cells. We reasoned that incorrectly paired heavy and light chains would show a decreased correlation in their SHM frequencies relative to correctly paired chains. Thus, we computed the Pearson correlation coefficient between IGHV and IGK/LV mutation frequencies for cells expressing non-majority light chain VJL combinations from misclustered clones. We found no significant difference (p=0.926, Fisher’s r-to-z transformation and z-test) between the levels of correlation for misclustered clones (0.761) and for accurate clones (0.769), suggesting that erroneous barcoding was unlikely a concern. In addition, we considered the possibility of poor germline V/J gene annotation for the light chains leading to a false appearance of light chain inconsistency. Since higher SHM frequencies are associated with increasing V(D)J annotation errors, we compared the light chain mutation frequency in misclustered and accurate clones. We found that the average IGK/LV mutation frequency across cells expressing non-majority light chain VJL combinations in misclustered clones was not significantly higher than that across cells in accurate clones (p=0.999; **Supplemental Fig. 2B**). Overall, these results suggest that the observed confidence for accurately identifying clones using heavy chain-based clonal clustering was not deflated by a false appearance of light chain inconsistency created by erroneous barcoding or incorrect light chain germline gene annotation.

### Characteristics of misclustered clones suggest room for improvement in heavy chain-based clustering

Current distance-based, heavy chain-based clonal clustering methods utilize information confined to VJL combination and distances between junction sequences to identify clonally-related sequences. We investigated the characteristics of the heavy chains of cells from misclustered clones to determine whether there was additional information in the heavy chains that could improve the clustering. Gupta et al. noted that shorter heavy chain junction lengths rendered lower and possibly insufficient diversity for effectively distinguishing clonal members from non-clonal ones (13). However, we found no significant difference between the heavy chain junction lengths of accurate clones and those of misclustered clones (p=0.486; **Figure 2A**). Thus, it is unlikely that heavy chain junction length could serve as an effective indicator for misclustered clones.

**Figure 2.**
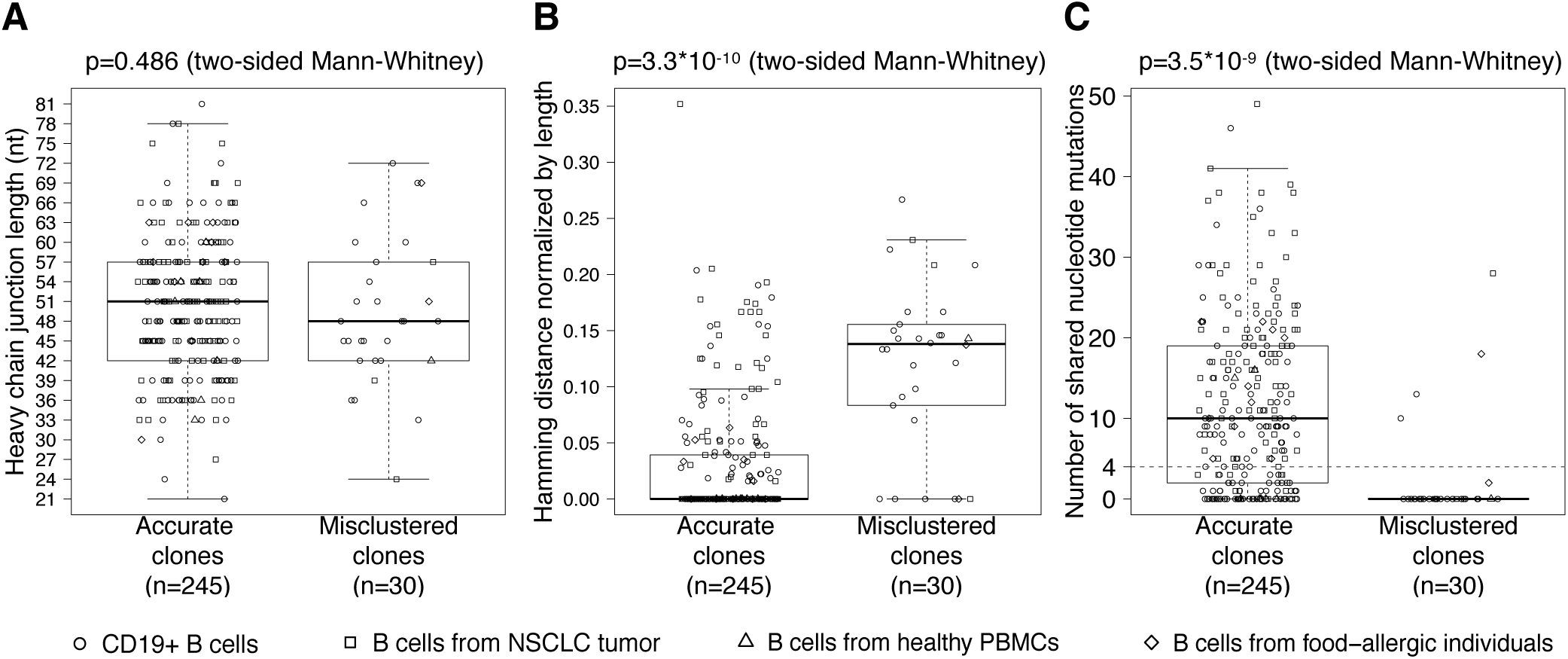
Characteristics of misclustered clones. **(A)** Heavy chain junction lengths of accurate clones versus misclustered clones. **(B)** The maximum pairwise distance between heavy chain junction sequences in accurate clones, versus the minimum pairwise distance between cells expressing inconsistent light chains in misclustered clones. **(C)** The minimum pairwise shared IGHV mutations in accurate clones, versus the maximum pairwise shared IGHV mutations between cells expressing inconsistent light chains in misclustered clones.

A central component of distance-based clonal clustering is the choice of a distance threshold that determines how dissimilar the junction sequence can be before it is unlikely for cells to be clonally related. To determine whether a better choice of distance threshold had the potential to correct misclustered clones, we compared the maximum pairwise distance between heavy chain junction sequences of cells in accurate clones, with the minimum pairwise distance between cells carrying light chains with different VJL combinations in misclustered clones. Cells with inconsistent light chains in misclustered clones had significantly more dissimilar heavy chain junction sequences compared with cells in accurate clones (p=3.3*10^-10^; **Figure 2B**). This implies that some of the misclustered clones could have been corrected by using a numerically lower (stricter) distance threshold, and that this lower threshold would not break apart the accurate clones.

True B cell clones are expected to share mutations resulting from SHM followed by clonal expansion and/or positive selection (12, 14). Hershberg & Prak suggested a minimum threshold of four shared mutations for inferring clones (12). We investigated whether cells in misclustered clones shared fewer mutations in their IGHVs compared to cells in accurate clones. To do so, we compared the minimum number of shared mutations between cells in accurate clones, with the maximum number of shared mutations between cells carrying light chains with different VJL combinations in misclustered clones. We found that cells carrying inconsistent light chains in misclustered clones shared significantly fewer mutations compared with cells in accurate clones (p=3.5*10^-9^; **Figure 2C**). Using Hershberg & Prak’s threshold of four shared mutations (12), 26 of the 30 misclustered clones (87%) would not have been clustered together based on their heavy chains (thus increasing specificity). On the other hand, within 4 of these misclustered clones, a subset of cells with consistent light chains would also become separated (thus reducing sensitivity). While the tradeoff between sensitivity and specificity needs further investigation, these results suggest that the extent of shared mutations in the IGHV is a potentially useful characteristic to consider in distance-based clonal clustering methods.

### Light chain information is insufficient for refining heavy chain-based clones

Given the availability of paired light chains in the single-cell datasets that we analyzed, we assessed the value added from that information. For misclustered clones (**Supplemental Fig. 2A**), this was trivial as any light chain inconsistency was immediately resolved by regrouping the cells into smaller clusters based on their light chain VJL combinations. For accurate clones, while the cells express consistent light chains, it is possible that these clusters may still contain multiple true clones grouped together. We thus investigated the extent to which further clustering cells based on the similarity of their light chain junctions (analogous to heavy chain-based clustering) would further split the accurate clones. When clustering using the heavy chains, a threshold around 0.2 normalized Hamming distance tended to separate clonally related cells from unrelated ones (**Supplemental Fig. 1C**). However, applying this same threshold to cluster light chains within accurate clones added virtually no information. The light chain junction regions of cells in accurate clones were highly similar and significantly more so compared with their heavy chain counterparts (Bonferroni-corrected p’s<0.001) (**Supplemental Fig. 2C**). In all but four of the accurate clones, the light chain junction sequence of a clonal member was at most 0.2 normalized Hamming distance away from the junction sequence of another clonal member most similar to itself (**Supplemental Fig. 2C**). In other words, clonal members tended to carry light chains with junction sequences that were at least 80% similar to each other.

We next investigated whether there would be further clustering based on light chain junctions at lower distance thresholds ranging, in increasing order of stringency, from 0.15 to 0.05 normalized Hamming distance, while bearing in mind that one could always artificially yield further clustering by imposing an increasingly stricter clustering threshold. At each clustering threshold, we determined the percentage of heavy chain-based clones that were further clustered on the basis of distances between their light chain junction sequences (**Table 2**). On average, 5% of the heavy chain-based accurate clones inferred via spectral clustering were further clustered at 0.15, the most lenient threshold explored. Even at 0.05, the strictest threshold explored, only 23.2% of the accurate clones were further clustered. This threshold is approaching the mean light chain SHM frequency, which ranges from 0.01 to 0.05 across the datasets, raising questions as to whether such further clustering is artificial rather than biological. Overall, light chain information does not support clonal clustering with greater granularity for the majority of heavy chain-based accurate clones.

**Table 2:** Percentage of accurate heavy chain-based clones further clustered based on light chains.

| *Clustering threshold for light chain junctions* | *0.05* | *0.10* | *0.15* |
| --- | --- | --- | --- |
| Dataset | Percentage of heavy chain-based clones<br>undergoing further clustering |  |  |
| CD19+ B cells | 20.3 | 6.8 | 3.0 |
| B cells from NSCLC tumor | 32.6 | 18.9 | 16.8 |
| B cells from food-allergic individuals | 40.0 | 10.0 | 0.0 |
| B cells from healthy PBMCs | 0.0 | 0.0 | 0.0 |
| Average | 23.2 | 8.9 | 5.0 |

## Discussion

In this study, we investigated the accuracy of heavy chain-based clonal inference. With single-cell VH:VL paired BCR datasets, we performed B cell clonal inference using only the heavy chains, effectively treating the datasets as if bulk-sequenced and unpaired. Over 80% of the inferred clones were accurate as defined by light chain consistency. Within the majority of these accurate clones, an additional round of clustering using the light chain sequence failed to yield finer resolution (<10% at a threshold of 0.1 normalized Hamming distance). Including a requirement that members of a clone share mutations between heavy chain sequences would have corrected 87% of the misclustered clones, though at the expense of also breaking up the accurately clustered part in 13% of these misclustered clones. Overall, while there remains additional information from the heavy chain that could be leveraged for improvement, we found heavy-chain based clustering alone capable of identifying clonal relationships with reasonable confidence.

Cells from four of the misclustered clones (13%) carried closely related heavy chains sharing a reasonable number of mutations in the V segment, yet they expressed light chains with different VJL combinations. There are several possibilities for how such clones could arise. During B cell maturation, the heavy chain rearranges first and the cell proliferates before the light chain rearranges (1). Thus, it is possible that the misclustered clonal relationships we detect represent daughter cells of the same heavy chain VDJ-rearranged B cell that developed different light chains. Alternatively, a group of identical, autoreactive, immature B cells could have undergone receptor editing, during which independent rearrangements gave rise to different light chains paired with an identical heavy chain rearrangement. In these scenarios, the heavy chain-based clonal clustering accurately reflects the underlying biology. However, we believe that these scenarios are unlikely as the chance that cells with identical heavy chains but different light chains would developmentally share the same temporal and/or spatial trajectories, in addition to being sampled together for sequencing, is expected to be very small.

When examining light chain consistency of cells from an inferred clone, if more than one light chain sequence was associated with a cell, we checked every sequence present for match against the clonal majority VJL combination. We did not, however, consider the possible complication where there was partial sampling in such a cell. Hypothetically, should two clonally related B cells with dual light chains each have a different light chain sequenced, our analysis would have considered a heavy chain-based clone containing these cells misclustered. However, such B cells have been reported to be rare, especially outside autoimmunity, with dual-κ and dual-λ cells estimated to occupy about 2-10% (24, 25) and 0.2% (26) of the normal murine repertoire, consistent with the percentages of cells with multiple light chains observed in the single-cell datasets here (1.1%-7.4%; all cells from the Croote et al. dataset had exactly one light chain).

Our results suggest several ways that heavy chain-based clonal clustering could be improved. For example, heavy chain junctions were more similar to each other in accurate clones than in misclustered clones, suggesting that the clustering threshold was perhaps too lenient for the latter. We also observed that accurate clones shared more mutations in their V segment. Some likelihood-based methods, such as *partis*, implicitly take into consideration shared mutations via the use of a multi-Hidden Markov Model that simultaneously emits multiple sequences (10). While computationally slower compared to the distance-based methods explored here, future studies should explore the potential benefit of such methods that use both junction distance and shared mutation patterns.

A limitation of this study is that all but one of the datasets contained relatively small numbers of B cells, and none was sorted for specific subsets, such as memory B cells, that would enrich for expanded clones. As a result, only a small number of the inferred clones contained multiple cells, and were thus suitable for analysis. As high-quality, single-cell B cell datasets of higher throughput with sorting for relevant B cell subsets emerge, similar analyses could be performed, leading to better estimates of performance.

Clonal relationship inference is an early step crucial for computational BCR repertoire analyses. As studies taking advantage of the relatively low cost of bulk BCR sequencing continue to generate unpaired BCR data, current clonal clustering methods can determine most clonal relationships with reasonable confidence using heavy chains only, and their performance may continue to improve by considering additional characteristics such as the number of shared mutations in the heavy chains.

## Acknowledgments

The authors thank Dr. Nima Nouri for critical reading of this manuscript.

