## Supplemental Data for "Immunoglobulin heavy chains are sufficient to determine most B cell clonal relationships^1^"

**A**

**Cell counts for 10x Genomics datasets at various stages of data processing**

| *Dataset Description* | *Platform* | *Source* | *Number of Cells* *Total* | *BCR contig(s) available* | *After filtering BCR contigs* | *After filtering for cells w/ 1 VH and ≥1 VL* | *After filtering for cells w/ transcriptomic profile of a B cell* |
|---|---|---|---|---|---|---|---|
| PBMCs (unsorted) of a healthy donor | V(D)J + 5' Gene Expression | AllCells (Catalog no. PB001) | 7726 | 1321 | 1255 | 1196 | 1115 |
| Cells (unsorted) from a squamous non-small cell lung carcinoma (NSCLC) tumor | V(D)J + 5' Gene Expression | A fresh surgical resection | 7802 | 2953 | 1817 | 1530 | 1388 |
| CD19+ B cells isolated from PBMCs of a healthy donor | Direct Ig Enrichment | AllCells (Catalog no. PB010-0) | 9465 | 9465 | 9219 | 8454 | - |
| B-lymphoblastoid cell line GM12878 | Direct Ig Enrichment | Coriell (Catalog no. GM12878) | 854 | 854 | 819 | 799 | - |

**B**

**Cell counts for Croote et al. 2018 dataset**

| *Subject* | *Number of B cells* *As in Croote et al., 2018* | *After filtering BCR contigs* | *After filtering for cells w/ 1 VH and ≥1 VL* |
|---|---|---|---|
| PA11 | 59 | 59 | 59 |
| PA12 | 267 | 265 | 265 |
| PA13 | 202 | 200 | 200 |
| PA14 | 212 | 211 | 211 |
| PA15 | 110 | 110 | 110 |
| PA16 | 123 | 123 | 123 |
| Total | 971 | 968 | 968 |

**Supplemental Figure 1. A.** Summary of datasets from 10x Genomics. **B.** Summary of B cells from food-allergic individuals. **C.** Distance-to-nearest neighbor plot to guide choosing a fixed distance threshold for inferring clones via hierarchical clustering. Cells were first partitioned into groups sharing the same IGHV gene, IGHJ gene, and heavy chain junction length. Within each partition, for each heavy chain junction sequence, the smallest non-zero nucleotide Hamming distance to other heavy chain junction sequences in the same partition was calculated. A threshold, indicated by a dashed line, was chosen via manual inspection to separate the bimodal distribution representing cells that were likely to be clonally related from unrelated ones.

**C**

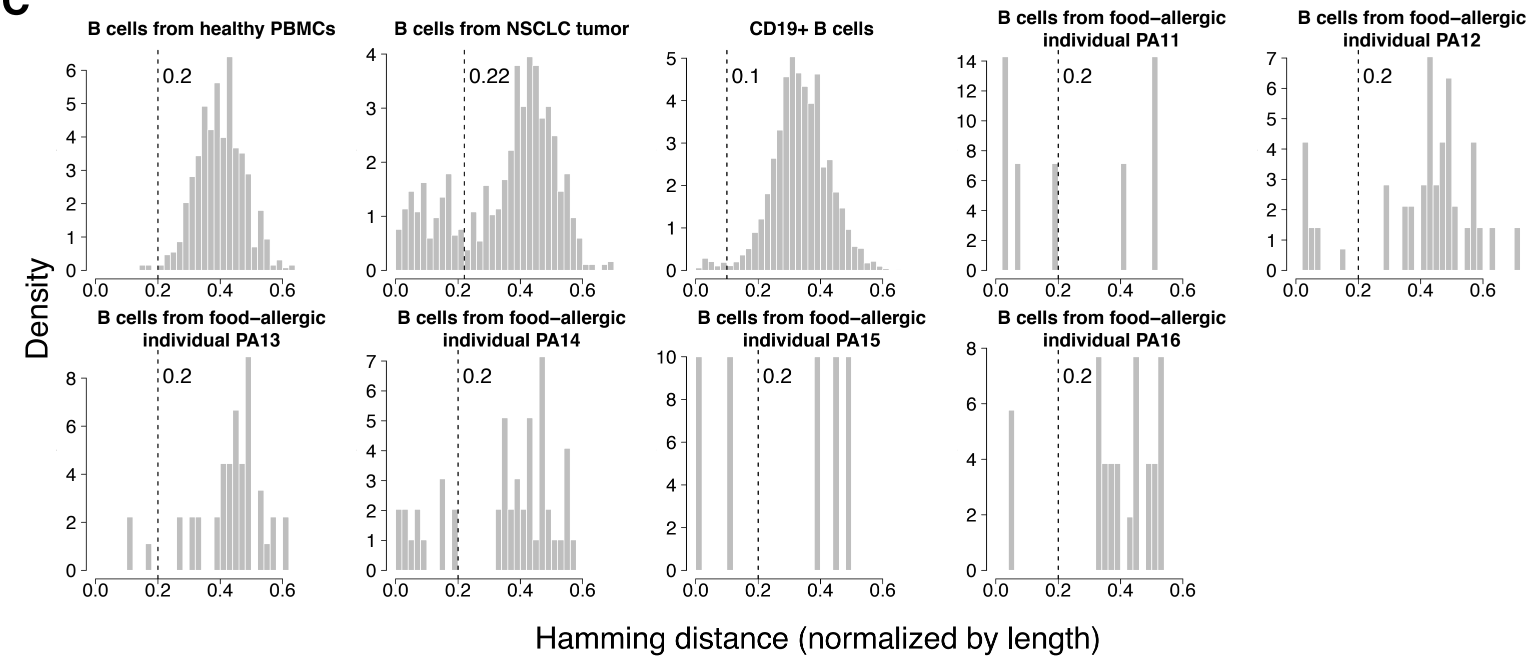

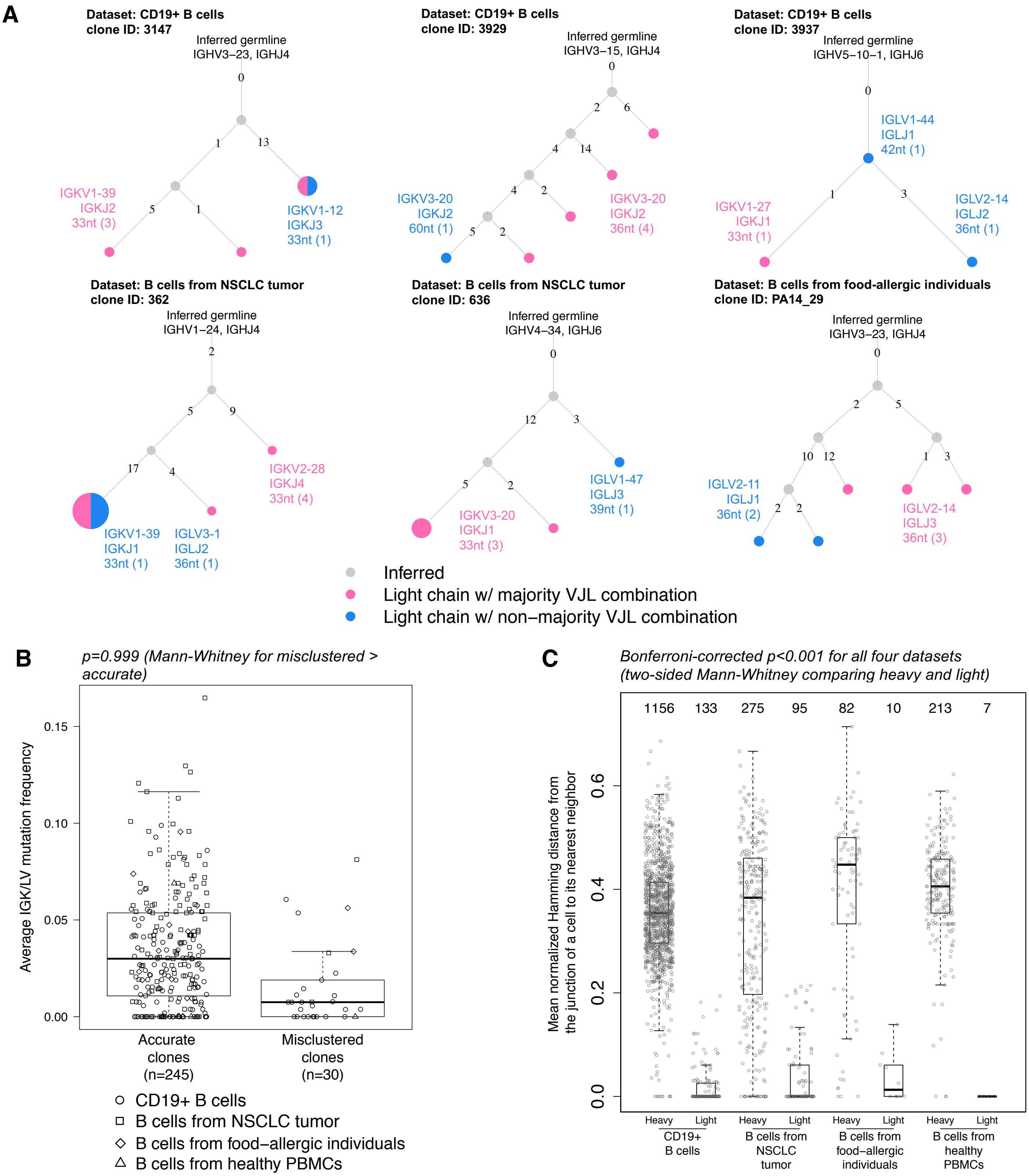


**Supplemental Figure 2. A.** Maximum-parsimony trees based on IMGT-numbered nucleotides 1-312 of IGHV for misclustered clones with >2 cells. Clonal members are shown as nodes and colored according to whether their light chain carries the clonal majority VJL combination. In case of a tie, a majority is designated arbitrarily (such as in Clone 3937). Cells with identical IGHV positions 1-312 share the same node, with node size proportionate to the number of cells at that node. Edge labels represent the number of mutations accumulated from one node to the next. **B.** Average IGK/LV mutation frequency across cells in accurate clones compared with that across cells expressing clonal non-majority VJL combination in misclustered clones. **C.** Means of the distributions of the smallest normalized Hamming distance between heavy chain junctions in partitions of cells with the same heavy chain VJL combinations, and between light chain junctions in accurate clones inferred via spectral clustering. A distribution was calculated for each heavy chain-based partition or clone. The mean of each distribution is visualized.
